# Sexual dimorphisms in phrenic long-term facilitation following severe acute intermittent hypoxia

**DOI:** 10.1101/2025.08.25.672148

**Authors:** Brendan J. Dougherty, Jessica M.L. Grittner

## Abstract

Rigorous pre-clinical research in male rodents defined the cellular mechanisms of respiratory neuroplasticity following brief exposures to hypoxia (acute, intermittent hypoxia; AIH). AIH elicits phrenic long-term facilitation (pLTF), a progressive increase in phrenic nerve amplitude over time. Mechanisms to AIH-induced pLTF are complex and variable depending on the severity of hypoxemia during AIH. Moderate AIH (mAIH; PaO_2_ ∼35-45mmHg) triggers spinal serotonin receptor activation to induce pLTF expression. More severe AIH (sAIH; PaO_2_ ∼25-30mmHg) induces pLTF through an adenosine receptor-dependent pathway. Here we assessed: 1) if sAIH-induced pLTF is expressed in female rats, and whether sAIH-pLTF is impacted by the estrous cycle; 2) if the magnitude of sAIH-induced pLTF in female rats is similar to male rats; and 3) whether GDX alters the magnitude of sAIH-induced pLTF. We hypothesized that female rats would express sAIH-induced pLTF, and that circulating steroid hormone levels would have minimal impact on sAIH-induced pLTF in either sex. Our findings reveal that female rats express robust pLTF (∼106% above baseline phrenic amplitudes) in response to sAIH, with minimal effects of estrous cycle stage. Female rats also showed a nearly 50% higher magnitude in sAIH-pLTF than males (p=0.006). Following GDX, pLTF magnitude was reduced in female rats (p=0.04), while males were unable to express pLTF. These findings predict unique cellular mechanisms to pLTF in female rats following sAIH, and sex-specific impacts of steroid hormone signaling on the expression of respiratory neuroplasticity.

## Introduction

Acute intermittent hypoxia (AIH) stimulates neuroplasticity in respiratory and non-respiratory motor circuits, revealing AIH as a therapeutic tool to augment task -specific neurorehabilitation^1–4^. Preclinical rodent studies over the past three decades have defined multiple cellular cascades giving rise to AIH-induced neuroplasticity in the respiratory motor system, with the dominant cellular mechanism being dictated by the severity of hypoxemia during AIH^3,5–7^. Clear modeling of this concept is observed with induction of phrenic long-term facilitation (pLTF), a specific form of AIH-induced respiratory neuroplasticity characterized as a progressive increase in phrenic neural output following brief episodes of hypoxia (i.e. AIH) in anesthetized, mechanically ventilated animals^8–10^. Two distinct, but related cellular pathways induce pLTF expression in this model: 1) a serotonin receptor-dependent pathway (the “Q-pathway”)^11–13^, and 2) an adenosine receptor-dependent pathway (the “S-pathway”)^14–16^. The Q-Pathway is the dominant mechanistic pathway to plasticity in response to moderate levels of AIH (mAIH; reducing PaO_2_ to ∼35-45mmHG), and the S-pathway dominates when more severe AIH doses are utilized (sAIH; PaO_2_ ∼25-30mmHg)^16–18^. Additionally, the S and Q-pathways regulate each other through mechanisms of cross-talk inhibition, adding complexity and redundancy to these pathways to neuroplasticity^19^.

Studies from our laboratory^20–22^ and others^23–25^ have demonstrated that sex hormones influence the development of mAIH-induced pLTF. In young-adult females, pLTF is only expressed during stages of the estrous cycle with high levels of circulating 17β-estradiol (E2)^20^, the most neuroactive form of estrogen. Removal of the ovaries (ovariectomy, OVX), the primary source of circulating E2, eliminates AIH-induced pLTF^20^. E2 supplementation - either systemically or locally to the vicinity of spinal phrenic motor neurons - restores the capacity for pLTF, providing strong evidence that E2 is a necessary factor to permit mAIH-induced neuroplasticity in females. In 2019, Hocker and colleagues demonstrated for the first time that female rats also express pLTF in response to sAIH^26^. Young-adult, ovariectomized (OVX) female rats, supplemented with physiological levels of exogenous E2, expressed pLTF following sAIH^26^. This finding demonstrated that females retain the capacity for pLTF across a range of AIH intensities under the right cellular conditions, and that supplemental E2 is sufficient to enable pLTF when serum E2 levels are reduced. The extent to which E2 preconditions are *necessary* for sAIH-induced pLTF via S-pathway mechanisms in gonadally-intact females is unknown. The first Aim of this study was to determine whether young-adult female rats with intact ovaries exhibit pLTF in response to sAIH across the normal estrus cycle, and whether this pathway is maintained following OVX. We hypothesized that sAIH would induce a similar magnitude of pLTF in *all* female rats, independent of estrus cycle stage or ovarian state, establishing that steroid hormone signaling is not prerequisite for sAIH-induced pLTF in female rats.

Male rats also rely on sex steroid hormones to express AIH-induced pLTF. The magnitude of mAIH-induced pLTF is reduced with age-related declines in circulating testosterone^23^, the principal male sex hormone, and removal of the testes, the primary source of circulating testosterone, eliminates the expression of pLTF in young-adult male rats^25^. Testosterone supplementation is sufficient to restore pLTF expression in castrated male rats supporting the idea that sex hormones are needed to permit respiratory neuroplasticity in both sexes. However, Zabka et al. demonstrated that the capacity of testosterone to restore pLTF following castration was reliant upon the enzymatic conversion of testosterone to E2 via aromatase; blocking aromatization eliminated the testosterone-induced rescue of mAIH-induced pLTF following castration^25^. These findings support the idea that E2 signaling is necessary in male rats, versus testosterone signaling, for induction of mAIH-induced pLTF following castration. How E2, or any other sex hormone signaling, permits the expression of sAIH-induced pLTF in gonadally-intact males is unknown. Therefore, the second Aim of this study was to assess whether male rats expressed sAIH-induced pLTF following removal of the testes. We hypothesized that *all* male rats would express severe AIH-induced pLTF regardless of gonadal status, establishing that sAIH-induced pLTF is also not reliant on sex hormone signaling in male rats.

## Methods

### Animals and groups

All experiments were performed with young-adult, Sprague-Dawley rats (3-5 months old; Envigo). The rats were housed in pairs in an AAALAC accredited vivarium with 12:12 hour light cycles and *ad libitum* food and water. The general term gonadectomy (GDX) is used for simplicity to denote removal of the testes in males, or removal of the ovaries in females. GDX in both sexes results in a significant and prolonged reduction in circulating steroid hormones^21^. Intact females were staged daily to determine estrus cycle stage and were studied either during the proestrus stage (high circulating estrogen levels) or estrus stage (low circulating estrogen levels). These stages were used to contrast findings from studies of moderate AIH (mAIH) where females demonstrate pLTF only during the proestrus phase and not the estrus phase^20^. In this study, rats were exposed to severe AIH (sAIH) or were time controls (i.e., no AIH exposure). The groups that were exposed to severe AIH (sAIH) were as follows: female, intact, proestrus (Pro., n = 5); female, intact, estrus (Est., n = 5), female post-GDX (GDX-female, n = 5), male intact (Intact-male, n = 6), male post-GDX (GDX-male, n =6). The time control groups matched the sAIH groups in terms of sex and gonad or hormone status.

### Estrus cycle staging

Female rats in the intact groups underwent daily vaginal cytology^20,22^. Vaginal cells were collected using autoclaved swabs soaked in distilled water and inspected under light microscopy. Estrus cycle stage was determined by agreement between two trained judges^22,27,28^. Based on staging, females were assigned to either the proestrus or estrus groups, and neurophysiology data collection was performed immediately after staging was completed.

### Gonadectomy surgery

All gonadectomy (GDX) surgeries were completed under aseptic conditions and are described in prior studies^21,29^. Animals were given subcutaneous, sustained-release buprenorphine 2 hours prior to surgery for pain (1 mg/kg, Wedgewood Connect). At the time of surgery, rats were sedated in a drop box with isoflurane and transferred to a nose cone with 2-5% isoflurane in 50-60% O_2_ (balance N2). Depth of sedation was confirmed by absence of toe pinch reflexes.

Females were placed prone, hair shaved from lumbar area of the back, and skin scrubbed with chlorhexidine. Bilateral incisions were made above the ovaries, in the dorsolateral region of the back. Ovaries were externalized and removed with a cautery. Remaining tissue (e.g., fallopian tubes, adipose tissue) was placed back into the body cavity, incised muscle was closed with absorbable sutures, and the skin secured with surgical clips. Males were placed supine, and the scrotum shaved and scrubbed with chlorhexidine. Bilateral incisions were made in the scrotum; the testicles were externalized and removed with a cautery. Vas deferens and connective tissues were placed back into the scrotal sack, and the incisions were closed with veterinary surgical glue (GLUture; Zoetis). At completion of the procedure, rats were removed from anesthesia and returned to their home cages. Rats were monitored for 72 hours for surgical complications or pain symptoms. Surgical clips were removed 7 days after surgery in females. Phrenic nerve recordings were completed 2 weeks following completion of GDX surgeries.

### Phrenic nerve recordings: surgical preparation

On the day of phrenic nerve recordings, rats were sedated in a closed chamber with isoflurane and then transferred to a custom-built warming table where isoflurane was maintained via nose cone (∼3% isoflurane in 50-60% O_2_, balance N2; Somnosuite, Kent Scientific). Tracheostomy and conversion to mechanical ventilation was completed (VentElite small animal vent, Harvard Apparatus), and bilateral vagotomy was performed. Isoflurane anesthesia was maintained until femoral arterial and venous catheters were placed, and then anesthesia was slowly converted to intravenous (i.v.) urethane (1.8 mg/kg). The arterial catheter was connected to a pressure transducer (SP844, MAMScap) for continuous blood pressure monitoring. Once the conversion to urethane was complete, a continuous i.v. infusion of 20% sodium bicarbonate in lactated Ringer’s solution was delivered at a rate of approximately 0.5 mL/hour to maintain blood pressure and acid-base balance throughout the surgery and data collection. Concentration of inspired O_2_ was monitored with an O_2_ sensor (AII 3000A, Analytical Industries) attached to the inspired line of the ventilator. Body temperature was monitored via rectal thermometer, and end-tidal CO_2_ was monitored with a flow-through capnograph (Respironics). These measurements as well as the nerve signals described below were synchronized using PowerLab 8/35 (AD Instruments). The left phrenic nerve was dissected via a dorsal approach. It was cut distally, desheathed, and placed on a bipolar silver hook electrode. A paralytic was administered (pancuronium bromide, 2.5 mg/kg) to remove extraneous movement artifacts during recordings. Nerve activity was amplified (gain, 10,000, A-M Systems) and bandpass filtered (300 Hz - 10 kHz). The resulting raw signal was recorded, rectified and integrated (time constant, 50 ms) by LabChart (version 8.1, AD Instruments).

### Phrenic nerve recordings: data collection and AIH protocol

Nerve recordings were not initiated until ∼45-60 min post urethane conversion to enable washout of isoflurane. Once stable nerve signals were established, the CO_2_ apneic and recruitment thresholds were established. The apneic threshold was found by progressively lowering the concentration of inhaled CO_2_ until phrenic bursting ceased. Once apnea was established, the inhaled CO_2_ was steadily increased in 1 mmHg increments to find the recruitment threshold (when phrenic neural bursting resumed). End-tidal CO_2_ was set to 2-3 mmHg above the recruitment threshold for the remainder of the experiment. A baseline blood sample was taken and analyzed for standard base excess, pH, PaCO_2_, and PaO_2_ (OPTI Gas). Animals were then exposed to sAIH consisting of 3 x 5-minute hypoxic episodes with a goal PaO_2_ of 40-55 mmHg^14,15^. Blood gas measurements were taken during hypoxic challenges to confirm hypoxia severity as well as at 15, 30, and 60 minutes after the last hypoxia. The time control animals had blood gas measurements taken at time points that were commensurate with the sAIH groups, though no hypoxia was administered.

Animals were excluded from the study if blood gas values fell outside of predetermined ranges and could not be corrected. Arterial PaCO_2_ was maintained isocapnic at ±1.5 mmHg baseline. At the end of the experiment, nerve signal integrity was confirmed with a hypercapnic-hypoxic challenge, and then all rats were euthanized by urethane overdose.

### Statistical analyses

Physiological data (e.g., weights, blood pH, PaCO_2_, PaO_2_, and base excess) were all compared using a 1-way ANOVA between same-sex groups. One-way ANOVA were used to compare the phrenic amplitude during hypoxia and the percentage change from baseline at 60-minutes after the final hypoxia. Two-way ANOVA with repeated measures was used to compare time versus group effects in phrenic amplitude percentage change from baseline at 15-, 30, and 60-minutes after the last hypoxia. Tukey’s multiple comparisons post hoc analyses were conducted when significance was found. Results were considered significant if p-values were less than or equal to 0.05.

## Results

Our first set of studies were designed to test the hypothesis that severe, acute intermittent-hypoxia (sAIH) would induce pLTF in female rats, independent of estrus cycle stage. The neural response to hypoxia and magnitude of sAIH-induced pLTF were compared in young-adult female rats in proestrus, notable for high circulating E2, and estrus (low E2)^20^. Physiological variables between experimental groups during baseline, hypoxic challenge, and 60 min post-hypoxia are provided in **Table 1**. One way ANOVA compared values across groups within each of these time domains. No significant differences were noted in core body temperature, or PaCO_2_; values were always similar between groups (**Table 1**). Though the means were not considered significantly different following ANOVA analysis (p=0.051), pH was slightly lower in estrus rats (7.378±0.01) relative to proestrus rats (7.422±0.01) during baseline which registered as significant with Tukey post -hoc analysis (p=0.042, **Table 1**). Lower pH during hypoxia in the estrus group (7.357±0.01) was also different from time control rats (7.404±0.01; p=0.044) during hypoxia (**Table 1**). Hypoxic challenge caused a reduction in PaO_2_ in experimental groups that was considerably lower than control rats. Specifically, proestrus (29.6±1.5 mmHg) and estrus rats (29.0±0.8 mmHg) similarly reduced PaO_2_ relative to each other (p=0.998), but this reduction was significantly lower than time controls during the same time period (223±11; p<0.0001; **Table 1**). This was fully expected since time control groups did not receive hypoxia. Likewise, though MAP was similar among groups during BL and at 60 min post-sAIH, hypoxia caused a significant reduction in MAP in proestrus (89±5 mmHg; p=0.041) and estrus (73±7 mmHg; p=0.002) rats compared to time controls rats (116±8 mmHg). Finally, a significant statistical difference in SBE was observed in proestrus rats (2.3±0.3 mequivl-1) compared to time control rats (−0.9±0.8 mequivl-1) during baseline (p=0.018), but both groups were within normal physiological limits (−3.0 and 3.0).

**Table 1.**
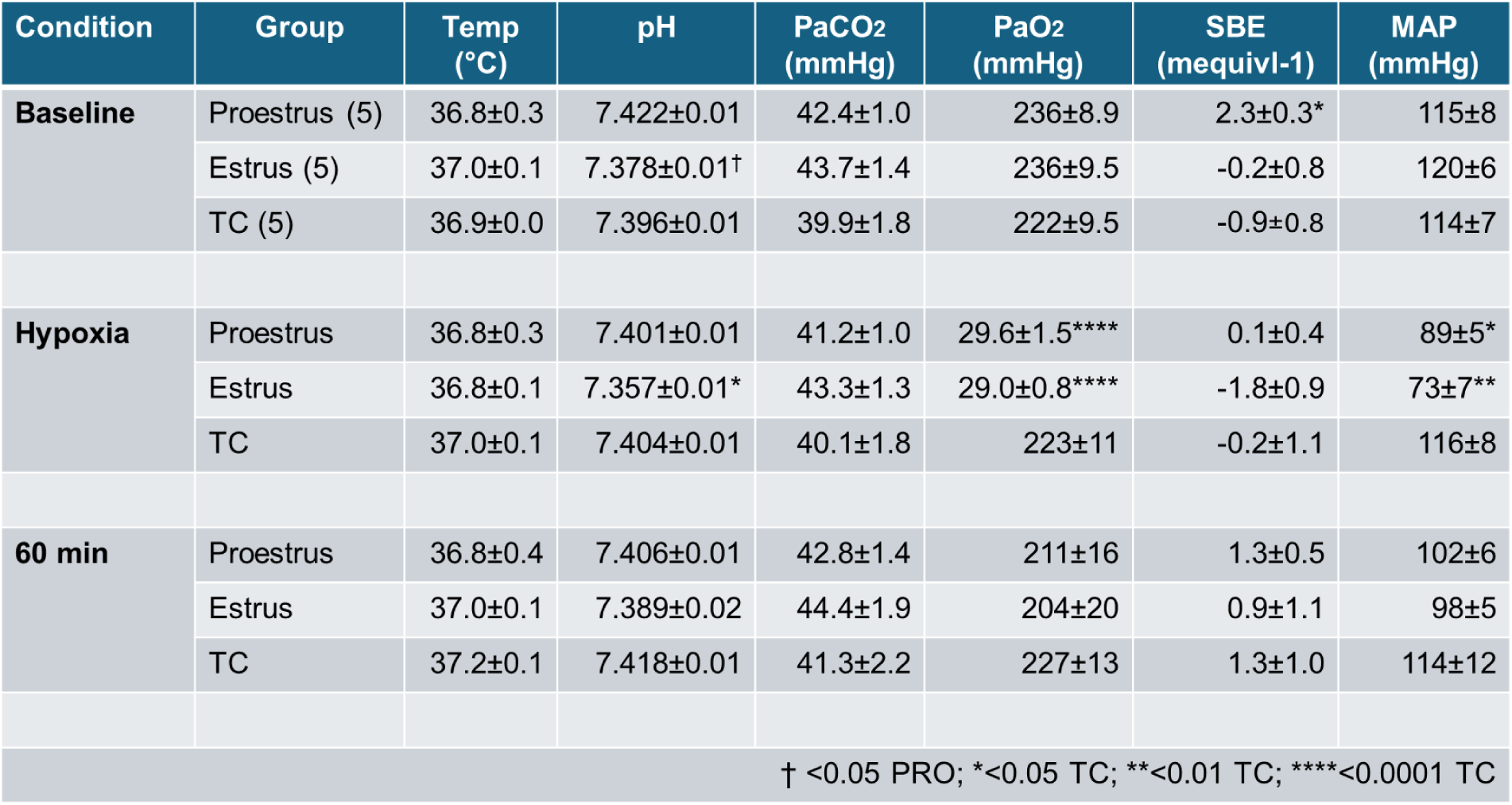
Physiological variables in female rats across the estrus cycle during sAIH-induced pLTF experiments.

**Figure 1A** displays the change in phrenic amplitude relative to baseline during hypoxic challenge in proestrus and estrus female rats compared to time control rats (no hypoxia). All female rats showed a robust neural response to hypoxia regardless of estrous cycle stage. This response was reflected in significantly elevated phrenic neural amplitudes relative to time control rats (**Fig. 1A**; p<0.0001). Phrenic amplitude in proestrus rats reached 208%±22% of baseline during hypoxia, while estrus rats showed a 163%±8% increase. These values were not statistically different from each other (p=0.091), which supports prior work demonstrating that hypoxic responses are not impacted by estrous cycle stage in females^21,30–32^. **Figure 1B** illustrates the phrenic neural responses as a percent change from baseline in the 60-min following sAIH between the three experimental groups. Two-way repeated measures ANOVA showed significant main effects of group (p<0.0001) and time (p<0.0001), and a significant group x time interaction (p=0.0003). By 30 min post-sAIH, both proestrus (85.7%±10% from BL; p=0.002) and estrus (70.4%±8% from BL; p=0.005) rats were significantly elevated above time control rats (8.99%±10% from BL) and above baseline values (p<0.004 for both; **Fig. 1B**). Phrenic amplitude continued to elevate above baseline values in proestrus and estrus rats, signifying robust sAIH-induced pLTF. By 60 min post-sAIH, proestrus (121.5%±18% from BL; p=0.004) and estrus (91%±14% from BL; p=0.004) rats were significantly elevated above time control rats (−1.14%±4% from BL) and above baseline values (p<0.01 for both; **Fig. 1B**). Importantly, at no time following sAIH were the changes in phrenic amplitudes different between proestrus and estrus rats (**Fig. 1B**). Therefore, for all subsequent analyses, the estrus and proestrus groups were combined into one experimental group denoted as “*intact female*” (n=10).

**Figure 1.**
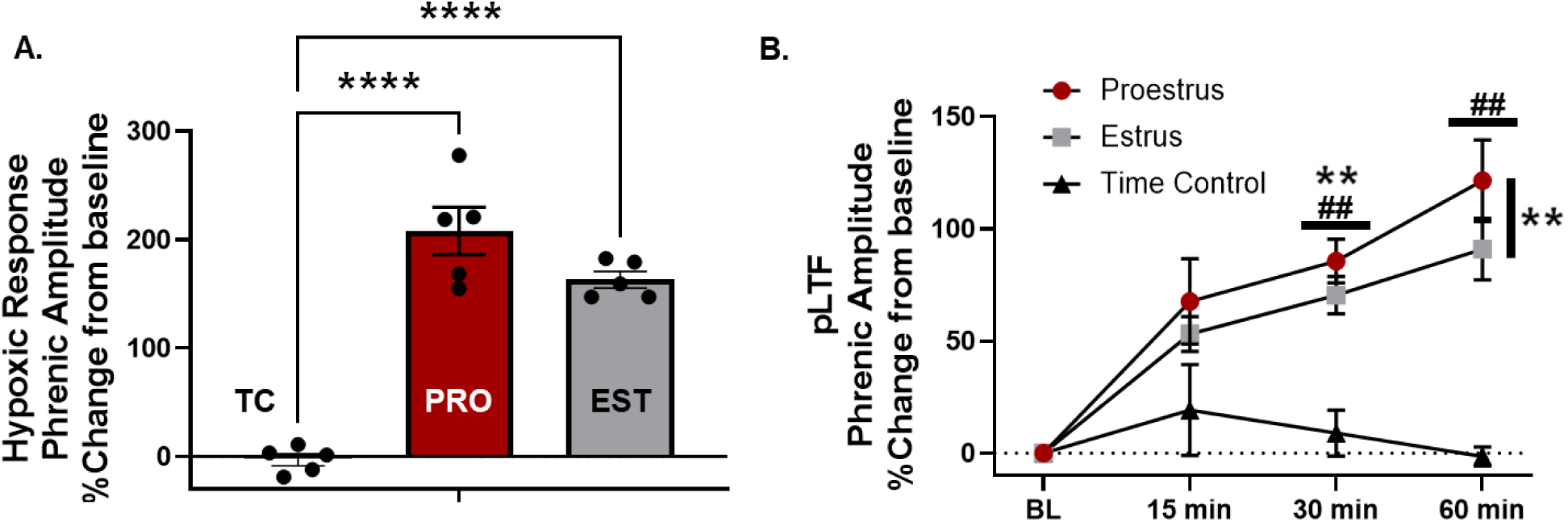
Hypoxic neural responses and sAIH-induced pLTF in female rats across the estrous cycle. Gonadally-intact female rats respond to sAIH, and express sAIH-induced pLTF across the estrous cycle. In (A.), female rats in proestrus (PRO; n=5) and estrus (EST; n=5) showed significantly elevated phrenic nerve amplitude in response to sAIH compared to time control rats (TC; n=5; no sAIH). Subsequently, in the 60 min following sAIH, female rats in proestrus and estrus showed a progressive elevation in phrenic nerve amplitude relative to baseline consistent with pLTF (B.). 30 min post sAIH, the phrenic nerve amplitude in both experimental groups was elevated above baseline and time control groups. This significant and progressive elevation in phrenic nerve amplitude persisted 60 min post-sAIH in both proestrus and estrus rats (B.). ^##^p<0.01 from baseline; **p<0.01 from time controls; ****p<0.0001 from time controls.

Next, we wanted to directly compare the neural response to sAIH and the magnitude of sAIH-induced pLTF between gonadally-intact male and female Sprague Dawley rats. Age-matched male rats underwent identical sAIH or time control protocols and data were compared to intact female rats. Male and female time control rats were statistically similar and thus combined into a single group (“*time control;”* n = 8). **Figure 2A** shows robust hypoxic responses from male (117.5%±17% from BL; p<0.0001) and female rats (185.7%±17% from BL; p<0.0001) that were significantly elevated relative to time control groups not receiving sAIH (1.09%±4% from BL). Additionally, the hypoxic response in female rats was significantly higher than male rats (p=0.003) signifying sex differences in the neural response to severe hypoxia (**Fig. 2A**).

**Figure 2.**
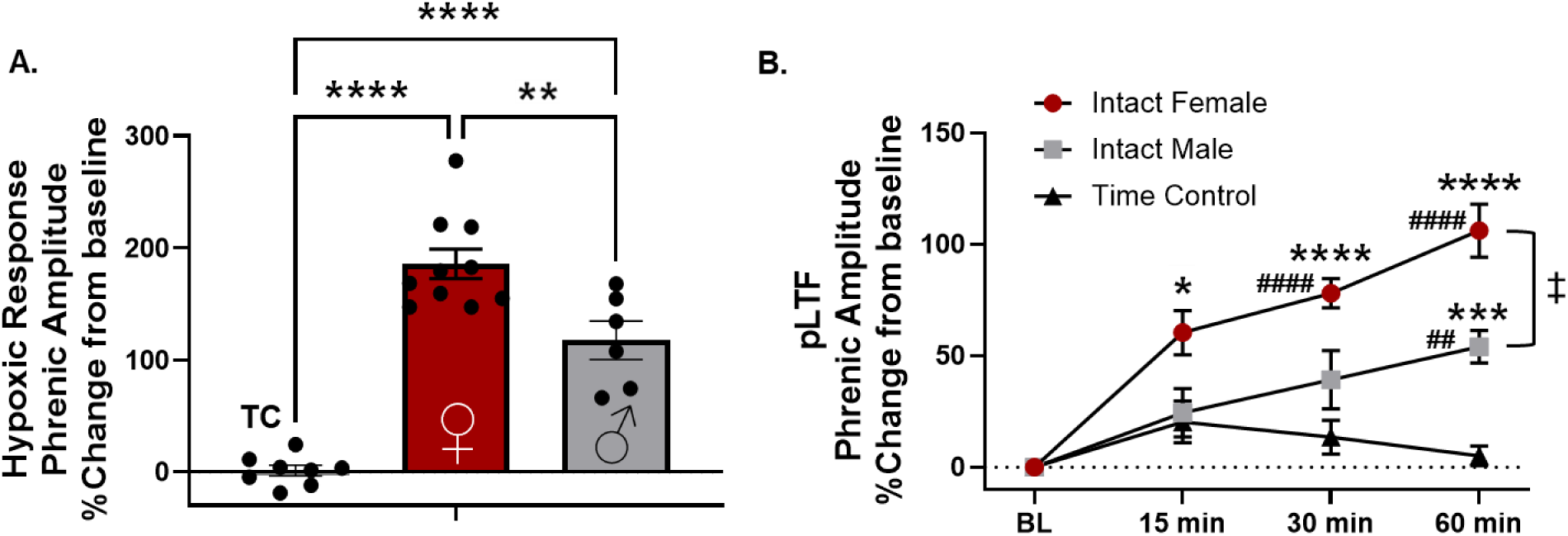
Hypoxic neural responses and sAIH-induced pLTF in gonadally-intact male and female rats. Female rats exhibit larger phrenic neural responses to severe hypoxia and larger sAIH-induced pLTF compared with age-matched males. The time control group is comprised of male and female rats. Both male and female rats showed a robust increase in phrenic neural output in response to sAIH (A.). The change in phrenic amplitude in response to sAIH in gonadally-intact female rats was significantly higher than age, and colony-matched gonadally-intact male rats (A.). Phrenic LTF was expressed in both sexes in response to sAIH as male and female rats showed a progressive increase in phrenic nerve amplitude relative to baseline and to time control rats (no sAIH stimulus) by 60 min post sAIH (B.). The magnitude of pLTF in females was significantly larger than males (B.) 60 min post-sAIH. ^##^p<0.01 from baseline; ^####^p<0.0001 from baseline. *p<0.05 from time control; ***p<0.001 from time control; ****p<0.0001 from time control. ^‡^p<0.01.

Assessment of sAIH-induced pLTF is shown in **Figure 2B**. Two-way repeated measures ANOVA revealed significant main effects for group (i.e. sex; p<0.0001) and time (p<0.0001) and a significant sex x time interaction effect (p<0.0001). Consistent with our first study, female rats showed significantly elevated phrenic amplitudes relative to time control rats following sAIH beginning at 15 min (60.4%±18% from BL; p=0.023), and extending to 30 min (78.0%±22% from BL; p<0.0001) and 60 min (106.2%±12% from BL; p<0.0001). These values were also significantly elevated from BL within the female rat group at all post-sAIH timepoints (p<0.001). Male rats also showed a gradual increase in phrenic amplitude resulting in pLTF by 60 min following sAIH. Male rats reached a mean phrenic amplitude of 54% from BL at 60 min which was significantly elevated from BL (p=0.003) and from time control (p=0.0009) groups ( **Fig. 2B**). Earlier male time points did not reach levels of significance. Therefore, by 60 min post-AIH, both male and female rats showed substantially elevated phrenic nerve amplitudes above BL and relative to time control rats signifying pLTF (**Fig. 2B**). The magnitude of pLTF was sexually dimorphic, however, as female rats had a significantly larger rise in phrenic amplitude (106.2%±12% from BL) compared to age-matched male rats (54%±7% from BL; p=0.006; **Fig. 2B**).

Finally, our third experiment explored the impact of gonadectomy (GDX) on sAIH-induced pLTF in female and male rats. GDX (removal of the primary reproductive organs; ovaries in females and testes in males) causes immediate and sustained reductions in circulating steroid hormone concentrations^21^. GDX female and male rats underwent identical sAIH protocols 2 weeks post-surgery to determine whether reduced steroid hormones impact the magnitude of the phrenic neural response to hypoxia or sAIH-induced pLTF. Data for the hypoxic responses in GDX rats were compared to rats from the intact female and intact male groups (study 2) and pLTF was assessed at 60 min post-sAIH (**Fig. 2B**).

GDX had minimal effect on the hypoxic response in female rats. With sAIH, the GDX female rats showed an average increase in phrenic amplitude of 217.1%±48% from BL, compared to 185.7%±17% for intact female rats (**Fig. 3A**; one way ANOVA; p=0.77). Both groups were significantly elevated relative to time control rats (−0.27%±5% from BL; p<0.0001; **Fig 3A**) who were not exposed to hypoxia. Similarly, GDX had minimal impact on the phrenic neural response to hypoxia in male rats. GDX male rats showed a mean increase of 128.2%±12% in phrenic amplitude from BL during sAIH, compared to 117.5%±17% increase for intact males (p=0.995). Like the female rats, both of male groups were significantly elevated above time control rats (p<0.001; **Fig. 3B**). Overall, the female rats had a larger response to hypoxia compared to the male rats. Though with the addition of the GDX groups, one way ANOVA confirmed only that GDX females remained statistically elevated compared with intact males (p=0.017) and GDX Males (p=0.040) while the intact female group was no longer significantly different from either intact (p=0.079) or GDX males (p=0.184; **Fig. 3A**).

**Figure 3.**
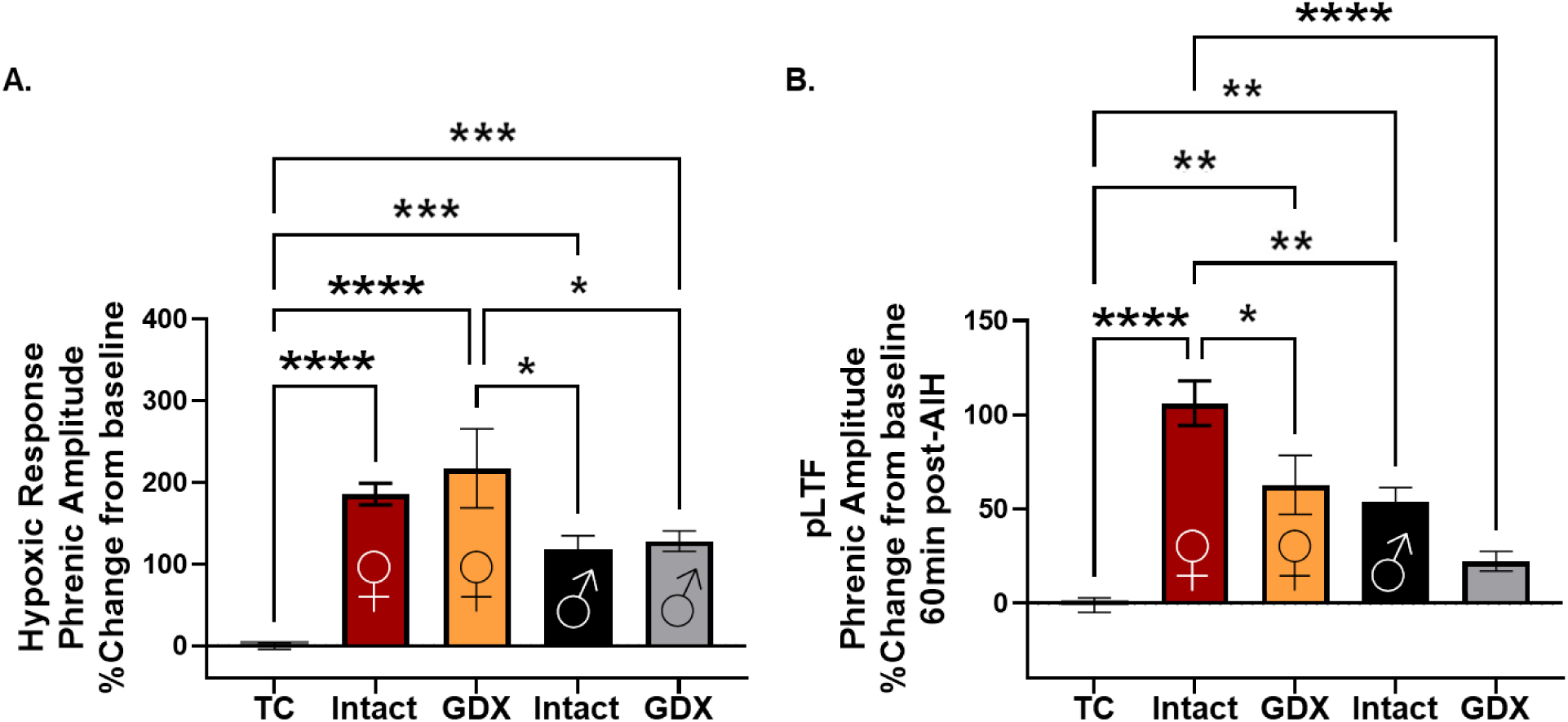
Effect of gonadectomy on hypoxic neural responses and sAIH-induced pLTF in male and female rats. Gonadectomy (GDX) resulted in sex-specific changes to the magnitude of sAIH-induced pLTF. Removal of the gonads did not impact the neural responses to hypoxia in either sex (A.). However, female rats, particularly those with GDX, showed higher hypoxic responses than males (A.). 60 min post-sAIH, phrenic amplitude was reduced in GDX female rats, but remained significantly higher than time control rats, indicating that female rats retained the capacity for sAIH-induced pLTF following GDX (B.). The magnitude of pLTF in GDX female rats was similar to intact male rats. Male rats lost the expression of sAIH-induced pLTF following GDX; male GDX rats had statistically similar phrenic nerve amplitude as time control rats (B.). *p<0.05; **p<0.01; ***p<0.001; ****p<0.0001.

GDX had a significant and sex-specific impact on the magnitude of sAIH-induced pLTF (**Fig. 3B**). In females, GDX caused a 42% mean reduction in pLTF magnitude 60 min post sAIH relative to gonadally-intact female rats (p=0.036, **Fig. 3B**). Intact female rats showed a mean elevated phrenic amplitude of 126%±36 above BL (**Fig. 3B**), compared to a mean of 53%±14 following GDX. Despite the GDX-associated reduction in pLTF magnitude, both intact females and GDX females retained the expression of pLTF relative to time control rats (p=<0.0001 and p=0.001 respectively). Thus, removal of circulating steroid hormones reduced, but did not eliminate the development of sAIH-induced pLTF in female rats. Of note, the magnitude of pLTF in GDX females was equivalent to the magnitude of sAIH-induced pLTF in gonadally-intact male rats (54%±7; p=0.98).

In male rats, GDX effectively eliminated the expression of sAIH-induced pLTF. Gondally intact male rats expressed pLTF with a mean magnitude of 54%±7 above BL, which was statistically higher than time control rats (p=0.003). Removal of the gonads reduced mean pLTF magnitude to 22%±5, which was not statistically different from the time controls (6.8%±7%, p=0.45). Though this reduced pLTF magnitude in GDX males was not statistically different from intact males when included in the collective one-way ANOVA presented in **Figure 3B**, when compared to intact males via T-Test (p=0.006), the difference in pLTF magnitude shows a statistical reduction.

Physiological parameters for studies 2 and 3 are provided in **Table 2**. Nearly all physiological variables were similar between groups within each measured time domain. As expected, all experimental groups receiving sAIH showed significantly reduced PaO_2_ (p<0.0001 for all) and MAP (p<0.01for all) during hypoxia when compared to time control rats not receiving sAIH (**Table 2**). Also, SBE in the GDX males during hypoxia (−1.7±0.6 mequivl-1) was lower than time control rats (1.0±0.6 mequivl-1; p=0.026), though both were well within appropriate ranges (3 to −3 mequivl-1) and MAP in the GDX males (96±8 mmHg) was reduced relative to time control rats (120±6; p=0.041) 60 min post-sAIH.

**Table 2.**
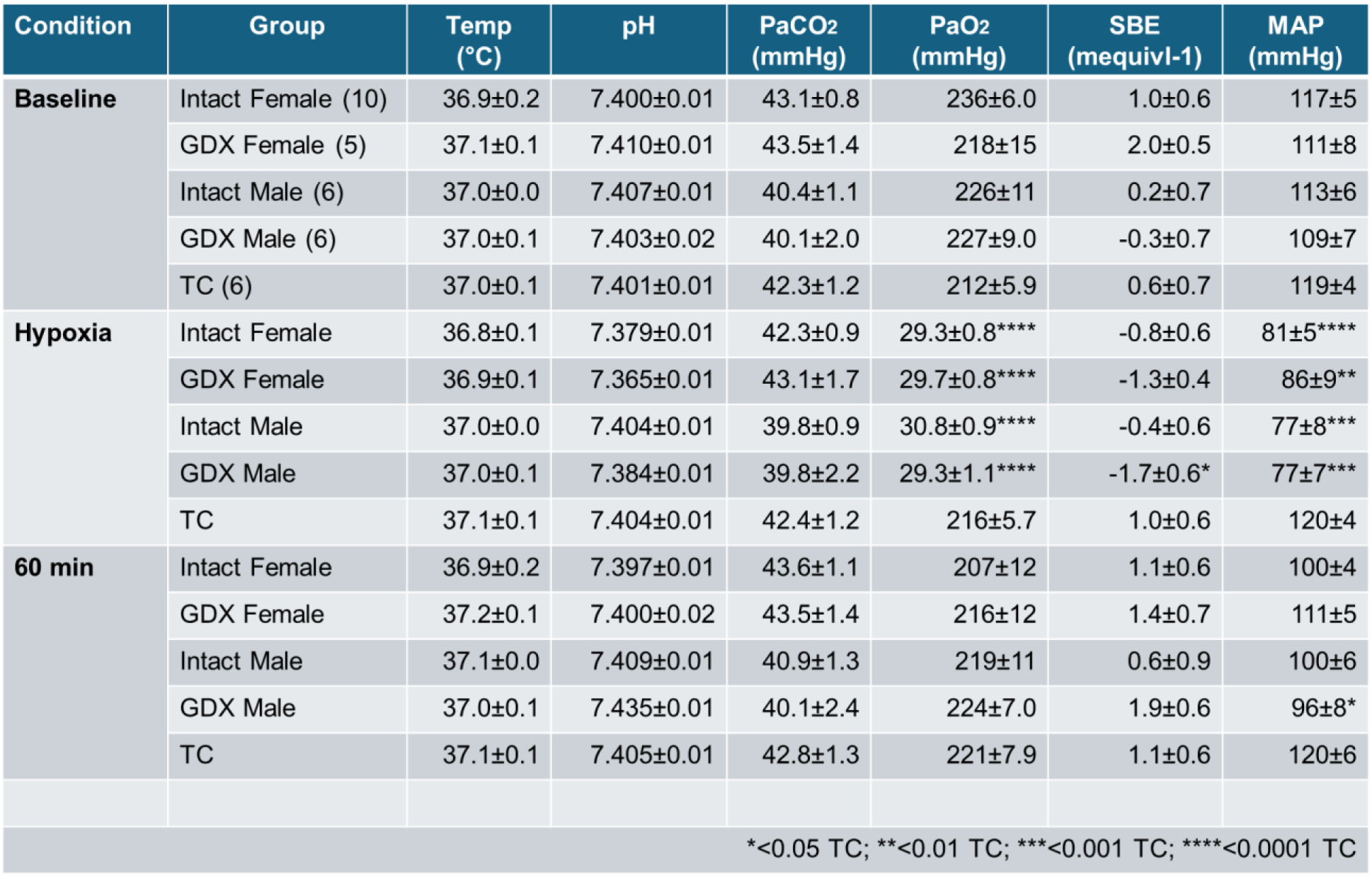
Physiological variables in male and female rats during sAIH-induced pLTF experiments.

## Discussion

This study had three primary goals: to evaluate the expression of phrenic long-term facilitation following severe acute-intermittent hypoxia in female rats across the estrous cycle, in female rats relative to male rats, and to determine the influence of circulating steroid hormones on the magnitude of this well-characterized form of respiratory neuroplasticity. Due to the unique mechanisms underlying sAIH-induced pLTF (relative to mAIH-induced pLTF)^14^ and prior studies demonstrating significant resilience in this pathway^33^, we hypothesized that females and males would express sAIH-induced pLTF of equal magnitudes, and that sAIH-induced pLTF would be minimally impacted by steroid hormones; pLTF would not fluctuate across the female estrous cycle, nor following GDX in either sex. Instead, our novel findings revealed sexual dimorphisms in the magnitude of sAIH-induced pLTF and sex-specific effects of GDX. In specific, young-adult, gonadally-intact female rats expressed robust pLTF in response to sAIH that was conserved across the estrous cycle, suggesting that normal fluctuations in circulating steroid hormones had minimal impact on this form of respiratory neuroplasticity in young-adult females. However, contrary to our hypotheses, the magnitude of pLTF in female rats was significantly higher than in male rats and removal of the primary reproductive organs (i.e. GDX) produced sexually dimorphic effects. GDX led to loss of sAIH-induced pLTF in males, and a reduction, but not loss, of pLTF in females. Collectively, these findings demonstrate that circulating sex steroids play unique, and sex-specific roles in sAIH mechanisms to pLTF.

### Female rats express robust sAIH-induced pLTF across the estrous cycle

Hocker and colleagues were first to demonstrate sAIH-induced pLTF in female rats in a study using ovariectomized female rats supplemented with physiological doses of E2^26^. Here we demonstrate for the first time that young-adult, *gonadally-intact female rats* express significant pLTF in response to sAIH. The magnitude of pLTF did not substantially differ across stages of the estrous cycle representing high serum E2 (proestrus) and low serum E2 (estrus), indicating that normal fluctuations in sex steroid hormones, E2 in particular, does not impact sAIH-induced pLTF. Thus, our findings are in agreement with Hocker and colleagues. Collectively, evidence now clearly demonstrates that female rats retain the capacity for respiratory neuroplasticity following sAIH. These findings validate the notion that the respiratory motor system in females maintains a similar ability to adapt across a wide range of physiological stimuli through expression of neuroplasticity as males.

### Females express greater hypoxic response and higher magnitudes of sAIH-induced pLTF than males

In order to provide sex-specific context to our results we directly compared the neural response to severe hypoxia and the magnitude of pLTF in females relative to age-matched males. Gonadally-intact female rats produced nearly 50% higher hypoxic responses compared to intact male rats. This outcome is consistent with several prior studies and demonstrates that female rats exhibit greater acute hypoxic phrenic responses compared to males^26,34,35^. The magnitude of pLTF 60 min post sAIH was also significantly higher in gonadally-intact females than in males; an unexpected result. With this finding, our results diverged from Hocker and colleagues who showed statistically similar magnitudes in sAIH-induced pLTF between males and ovariectomized females supplemented with E2^26^. Yet, a closer comparison of the data shows pLTF outcomes that are more similar than expected between the studies. For example, the magnitude of sAIH-induced pLTF in males (61%±69 vs. 54%±7) and females (130%±22 vs. 106.2%±12%) were similar, but higher variability in the findings of Hocker and colleagues likely masked sex -differences^26^. Regardless, our findings showed a clearly elevated pLTF in gonadally-intact female rats relative to age, colony, and vendor-matched male rats.

Whether the enhancement of pLTF magnitude is the result of unique, female-specific mechanisms or simply a more robust physiological response to the same cellular mechanisms is not yet known. In males, sAIH creates tissue-level hypoxia in the spinal cord, which triggers local microglia to release adenosine^7^. Subsequent activation of adenosine A2A receptors on phrenic motor neurons, and the activation of S-pathway mechanisms facilitate pLTF^15,36,37^. Downstream components of the S-pathway include activation of adenylyl cyclase/cyclic AMP signaling, and activation of exchange protein activated by cyclic AMP (EPAC)^38,39^, PI3 kinase/AKT^14,37^, mammalian target of rapamycin (mTORC1)^40^, the new synthesis of immature TrkB isoform^37^, and post-translational modifications of N-methyl-D-aspartate (NMDA) receptors^41,42^. Assuming S-pathway mechanisms are similar in female rats (which has not been confirmed), sAIH-induced tissue hypoxia may induce a greater microglial, or other glial cell, response and release of larger stores of local adenosine near phrenic motor neurons^7^. The subsequent activation of additional A2A receptors and downstream S-pathway signaling may enhance the magnitude of pLTF relative to males.

Alternatively, females may utilize unique, sex-specific pathways to sAIH-induced pLTF. Drug induced phrenic motor facilitation (pMF) studies were principally used to determine the cellular mechanisms of the S-pathway to pMF^18,37,43^. In one case, activation of 5HT-7 receptors with spinal application of AS-19 (5HT-7 receptor agonist), lead to a phenotypically similar facilitation of phrenic neural activity as induced with sAIH^43^, demonstrating one of several possible pathways to plasticity via the S-pathway. Though it is not believed that activation of 5HT-7 receptor is *necessary* for sAIH-induced pLTF in male rats *per se*, this, or similar receptors linked to the S-pathway^16,44,45^ may be coincidently activated in females, amplifying the impact of sAIH in a sex-specific manner.

Perhaps the most unanticipated results were the unique, sexually-dimorphic effects of GDX on sAIH-induced pLTF. GDX, and the associated loss of circulating steroid hormones, eliminated expression of sAIH-induced pLTF in male rats, demonstrating that these sex hormones have significant influence on the cellular pathways to all forms of AIH-induced pLTF. Male rats also lose the capacity for mAIH-induced pLTF when steroid hormone levels are reduced, either through GDX or natural aging^23,25^. The ubiquitous nature of these effects likely indicates that either: 1) steroid hormones work to modulate components of both S- and Q-pathways to pLTF simultaneously, or 2) that the effects of sex steroid signaling converge on cellular mechanisms that are necessary for both S- and Q-pathways. Recently, Nichols and Mitchell studied mechanisms of sAIH-induced pLTF and compared them with components of the S-pathway that were defined with drug activation (i.e. pMF studies)^14^. A unique finding was that sAIH-induced pLTF requires MEK/ERK activity, similar to mAIH-induced pLTF^46^. This result diverged from pMF findings demonstrating MEK/ERK signaling only in the Q-pathway to pLTF^46^. Thus, MEK/ERK may be common to both S- and Q-pathways to pLTF in males. Androgen receptors, activated by testosterone and dihydrotestosterone, are known to activate intracellular kinase cascades, including MEK/ERK^47^, via non-genomic signaling pathways^47,48^. Loss of testosterone following GDX may reduce basal activation of MEK/ERK and interrupt the S- and Q-pathways, limiting the expression of all forms of pLTF following AIH.

The MEK/ERK signaling pathway is also a downstream target of estrogen receptors^49–52^. In particular, activation of extranuclear estrogen receptors are associated with increased MEK/ERK^53^. Our group showed that activation of membrane-associated estrogen receptors using E2-BSA, a non-permeable estrogen conjugated to bovine serum albumin, was sufficient to restore mAIH-induced pLTF in female rats following GDX suggesting an important role for membrane estrogen receptors in permitting mAIH-induced pLTF^20^. Perhaps loss of circulating estrogen with GDX reduces membrane estrogen receptor activation and downstream MEK/ERK signaling, limiting the development of mAIH induced pLTF. If the MEK/ERK pathway is also involved with sAIH-induced pLTF in females similar to males^14^, then GDX would have similar downstream effects. However, since female rats retain pLTF following sAIH, albeit at reduced magnitude, estrogen receptor effects on MEK/ERK would not fully account for our results and would strengthen the argument for a female-specific mechanism to sAIH-induced pLTF.

### Technical considerations

One methodological consideration should be addressed to appropriately interpret our findings. Serum steroid hormone concentrations could not be confirmed in these studies which may have impacted our outcomes, specifically in study 1. Female rats experience a complete estrous cycle every 4-5 days separated into four clearly defined stages based on vaginal cytology^27,28^. These stages correspond to normal cyclic fluctuations in circulating steroid hormones. Proestrus is characterized by gradually increasing levels of circulating 17ꞵ-estradiol (E2), with peak E2 levels occurring in mid-late phase proestrus. E2 levels precipitously decline as female rats pass through the early portion of the subsequent estrus stage and reach the nadir of circulating E2 in the early-middle phase of estrus. Our prior work utilized these periods of naturally high E2 (proestrus) and low E2 (estrus) to demonstrate that high serum estrogen was necessary for the expression of pLTF following moderate AIH (35-45mmHG) in females^20^, different from the severe AIH (25-30mmHg) stimulus used here. Daily assessments of vaginal cytology were completed for a minimum of one week (or two full cycles) by at least two trained judges at a time, and rats were grouped according to the agreed upon cycle stage on the morning of pLTF experiments. In our prior work, grouping of rats in this manner was corroborated with serum E2 measurements^20^. In a few instances, rats were reassigned to either high or low E2 groups based on these follow -up measurements. Here, we were unable to confirm the grouping of our female rats into high and low E2 groups using a secondary serum E2 measurement due to a technical issue during ELISA assay processing. Though we are confident that our staging methods are accurate, fluctuations in E2 can occur quickly with rapid changes in cycle stages and, thus, there may be rats within the intact female groups that were not placed in the best grouping or fell within an intermediate range of serum E2, impacting our findings.

## Conclusions

Here we demonstrate that gonadally-intact female rats express phrenic long-term facilitation (pLTF) in response to sAIH regardless of estrous cycle stage. In addition, female rats show an enhanced magnitude of pLTF relative to gonadally-intact male rats, suggesting either a more pronounced activation of cellular pathways to sAIH-induced pLTF (i.e. the S-pathway), or a unique, female specific pathway producing a larger neuroplastic response. The concept of a female-specific mechanism is strengthened by data demonstrating sexually dimorphic impacts of GDX on sAIH-induced pLTF. Phrenic neuroplasticity is abolished following sAIH-induced pLTF in males, while female rats retain pLTF at levels similar to gonadally-intact male rats. Future studies will focus on deducing the unique underlying mechanisms of sAIH-induced pLTF in female rats.

Although severe AIH protocols are not considered translational given the potential side effect of such significant hypoxemia in humans, the interaction between the competing cellular mechanisms to AIH-induced plasticity offer potential targets to amplify translational benefits. For example, since S-pathway mechanisms are known to constraint the magnitude of mAIH-induced plasticity in males^7,18^, blocking this pathway prior to mAIH treatments may boost the clinical impact. Trumbower and colleagues demonstrated this idea by showing that caffeine, a common A2A receptor antagonist, augmented AIH-induced gains in walking function in individuals with chronic spinal cord injury^54^. Should there be unique, female-specific mechanisms to sAIH, then the chosen targets to enhance the clinical effects of AIH in women may be different. Our findings provide foundational data for which to discover these potentially clinically meaningful targets.

